# Matrix Rigidity Governs Switch-Like Oscillatory State Transitions of the Segmentation Clock in Isolated Presomitic Mesoderm Cells

**DOI:** 10.1101/2024.07.02.601712

**Authors:** Chun-Yen Sung, Yuqian Yang, Usha Kadiyala, Owen Blanchard, Liam Yourston, Derek Walker, Linyuan Li, Shiyu Sun, Zhaoyi Xu, Mariela Lopez Gonzalez, Jianping Fu, Qiong Yang

**Author notes:** These authors contributed equally to this work (Chun-Yen Sung, Yuqian Yang). These authors contributed equally to this work (Usha Kadiyala, Owen Blanchard, Liam Yourston).

## Abstract

The segmentation clock, a genetic oscillator in the presomitic mesoderm (PSM), is known to be influenced by biochemical signals, yet its potential regulation by mechanical cues remains unclear. The complex PSM microenvironment has made it challenging to isolate the effects of mechanical signals on clock behavior. Here we investigated how mechanical stimuli affect clock oscillations by culturing zebrafish PSM cells on bioengineered elastic substrates (PDMS micropost arrays) with tunable rigidities ranging from 0.6 to 1,200 kPa. We observed an inverse sigmoidal relationship between substrate rigidity and the percentage of oscillating PSM cells, with a switching rigidity threshold between 3-6 kPa. The oscillation periods of oscillating PSM cells showed a consistently broad distribution across the substrate rigidity conditions tested. Moreover, these oscillatory PSM cells exhibited distinct biophysical properties, including reduced motility, contractility, and sustained circularity, compared to non-oscillating ones. These findings highlight a role of cell-substrate interactions in regulating segmentation clock behavior, providing insights into the mechanobiology of somitogenesis.

**Highlights:**

- Oscillatory behaviors of single PSM cells respond to substrate rigidity in a switch-like manner, transitioning from an oscillatory state to a quiescent state at a critical rigidity threshold between 2.9 kPa and 6 kPa.
- Increased substrate rigidity significantly suppresses the percentage of oscillating PSM cells, while oscillation period and cycle number show no consistent rigidity-dependent trend.
- Non-oscillating PSM cells exhibit distinct biophysical properties compared to oscillating ones, including reduced circularity, a polarized and elongated morphology, greater motility, and increased contractility.
- Aggregates of PSM cells exhibit similar trends in response to substrate rigidity changes, except for increased oscillation percentages across different rigidity conditions, suggesting a potential interplay between cell-cell and cell-matrix communications in influencing clock oscillation behavior.

## Introduction

The rhythmic formation of somites during vertebrate embryogenesis is regulated by the segmentation clock, a genetic oscillator operating in the presomitic mesoderm (PSM) that relies on the periodic expression of cyclic genes involved in various signaling pathways, including the Hes/Her family, Delta/Notch, Wnt, and Fgf^1–4^. This clock exhibits notable spatiotemporal changes along the anterior-posterior (AP) axis of the PSM, including a progressive anterior slowing and eventual arrest of oscillations as cells incorporate into somites^5–8^, as well as a transition from desynchronized to synchronized oscillatory behavior among neighboring cells mediated by Delta-Notch signaling^9^. While biochemical regulation plays a crucial role in governing the segmentation clock’s dynamics, it alone is insufficient to fully explain these spatial variations in the segmentation clock’s behavior. Recent studies suggest that tissue mechanics may contribute to the regulation of the segmentation clock’s spatiotemporal properties^10–15^. In this study, we investigate how modulating mechanical forces changes the segmentation clock’s temporal properties, which remain largely unknown.

The mechanical properties of the PSM microenvironment undergo significant changes during somitogenesis. As mesodermal progenitor cells transition from the posterior to the anterior region of the PSM during zebrafish anteroposterior axis elongation, they encounter a stiffening process known as the “jamming transition”, which transforms the tissue from a fluid-like to a dryer foam-like architecture^13,14^. This transition is characterized by posterior to anterior spatiotemporal changes in the extracellular matrix (ECM) composition, cell density, and motility^10,14,16,17^. In the posterior PSM, the ECM primarily contains hyaluronic acid, while in the anterior region, the ECM becomes dense and stiff due to the increasing abundance of fibronectin and collagen fibers^15^. These observations suggest that in addition to the three-tier mode^1,18^ of the segmentation clock, which involves single-cell oscillators, cell-cell communications, and morphogen gradients, the mechanical gradient of the PSM microenvironment may act as a potential fourth tier of regulation that impacts the properties of the segmentation clock. Consistent with this idea, Hubaud et al. showed that dissociated mouse PSM cells respond to substrate adhesion property changes and switch between quiescent and oscillatory states via intracellular YAP-dependent regulation^11^. Nonetheless, these studies were conducted on glass surfaces, whose stiffness values are much greater than the physiological stiffness range of the PSM tissue. Thus, it remains unclear how PSM cell oscillators respond to varying mechanical stimuli within a physiologically relevant stiffness range, and how changes in cellular physical properties, such as cell morphology and migration, may correlate with the oscillatory behavior of individual PSM cells.

YAP has emerged as a key nuclear mechanotransducer that transmits mechanical signals from the extracellular environment to the cell nucleus. YAP translocation to the nucleus has been shown to be dependent on substrate rigidity, with stiffer or more adhesive substrates promoting nuclear localization^19^. In cultured mouse embryonic fibroblasts, YAP shows a Hill-like nuclear translocation response to substrate rigidity, with a reported rigidity threshold of approximately 5 kPa^20^. This mechanosensing mechanism is mediated by talin, a cytoskeletal protein that is involved in intracellular force transmission to the nucleus only above a threshold in substrate rigidity^20^. In some cellular contexts, YAP has also been reported to inhibit Notch in a cell-autonomous manner^21^. In the context of the segmentation clock, this raises the possibility that oscillatory behavior in individual PSM cells may depend on a YAP-dependent stiffness threshold. In mouse PSM cultures, Hubaud et al. further presented a co-regulation model for YAP and Notch signaling, in which YAP-dependent mechanical cues modulate an excitability threshold for oscillations *in vitro*, while Notch signaling, a well-established activator and synchronizer of the segmentation clock^22^, provides the stimulus that promotes these oscillations^11^. Studies in epidermal stem cells suggest a potential mechanistic link where mechano-activation of YAP/TAZ may indirectly regulate Notch signaling through transcriptional control of Delta-like ligand expression, acting as ‘in cis’ inhibitors of Notch^23^, although the molecular mechanisms connecting YAP and Notch to segmentation clock dynamics remain unclear.

Although single PSM cells function as self-autonomous oscillators with minimal cell-to-cell contact or juxtacrine Delta/Notch activity^4^, they may undergo YAP-mediated cis-inhibition of Notch due to mechanical interactions with the ECM. In contrast, cell aggregates within the PSM tissue may modulate the spatiotemporal features of the segmentation clock through the antagonistic interplay between trans-activation of Notch via Delta/Notch interactions among neighboring cells and YAP-mediated cis-inhibition of Notch via mechanical feedback.

In this study, we investigated how mechanical cues affect the oscillatory behavior of isolated and aggregated zebrafish PSM cells, dissociated from transgenic zebrafish embryos expressing cyclic Her1-Venus^5^, by culturing them on polydimethylsiloxane (PDMS) micropost arrays with tunable post spring constants. By varying the height and diameter of the microposts, the spring constant of PDMS microposts and thus the substrate rigidity could be precisely controlled, allowing us to investigate cellular responses to a range of mechanical environments physiologically comparable to PSM tissues^24^. This approach eliminates the influence of long-range morphogen gradients (Fgf, Wnt, and RA) present in vivo^8^, by isolating individual PSM cells or cell aggregates from their native tissue environment. As a result, cells are analyzed in the absence of spatial gradient inputs, enabling the examination of the intrinsic *her1* negative feedback oscillatory circuit at the single-cell level under defined mechanical conditions.

We report that the segmentation clock exhibits a switch-like response to changes in substrate rigidity, with a significantly reduced percentage of oscillating PSM cells above a stiffness threshold of 3.77 kPa, as determined by fitting a phenomenological model incorporating Hill-like YAP nuclear translocation. In contrast, the oscillation period, ranging widely from 60 to 100 minutes, shows no clear dependence on mechanical stimuli. This suggests that individual PSM cells may determine their period through an intrinsic pacemaker, likely driven primarily by transcriptional and translational delays in the *her1/7* negative feedback loop^25^ and influenced by other position-dependent biochemical signals. Consistent with findings from mouse PSM cultures^11^, our results in zebrafish PSM cultures suggest that mechanical inputs may act as a gating mechanism to determine whether cells remain oscillatory or become quiescent, potentially via regulation of intracellular YAP activity. Furthermore, compared to isolated cells, cell aggregates exhibit a higher probability of oscillations across all rigidity conditions tested in this study without a clear switching threshold, suggesting that the restoration of cellular interactions and tissue-level mechanics can co-modulate the segmentation clock dynamics of PSM cells. Thus, mechanical regulation of the segmentation clock in PSM cells could represent an additional tier of control, complementing the existing models based on genetic circuits, cell-cell communication, and morphogen gradients.

## Results

### Substrate Rigidity Modulates Single-Cell Segmentation Clock Oscillations

To examine the influence of substrate rigidity on the oscillatory behavior of isolated zebrafish PSM cells, we modified a zebrafish PSM single-cell protocol^4^ by using mechanical dissociation to minimize potential perturbations to cell morphology, cytoskeletal organization, and cell-matrix interactions associated with chemical dissociation. PSM cells were dissociated from tailbuds of zebrafish embryos at the 5- to 8-somite stage containing the *Tg(her1:her1-Venus)* transgene^5^, before being cultured on two distinct surfaces: Pluronic-coated glass coverslips, which inhibit cell-surface adhesion, and Matrigel-coated glass coverslips, which promote cell-surface adhesion (Figure 1A). Notably, cells isolated from the anterior PSM (A-PSM) cultured on Pluronic-coated glass exhibited earlier Her1-Venus oscillation peaks but only completed one cycle. In contrast, cells from the posterior PSM (P-PSM) initiated their oscillations later, but these oscillations lasted for multiple cycles, suggesting that the oscillation dynamics of individual PSM cells vary depending on their original location within the PSM, potentially indicating that cells retain positional information from their endogenous tissue environments (Figure S1A). This A-P axis spatial dependency observed in the oscillation cycle number is consistent with previous findings by Rohde et al.^8^. For the remaining results presented in this study, we exclusively utilized P-PSM cells given their capability of sustained oscillations. The Her1-Venus oscillatory behavior and morphology of single P-PSM cells displayed marked differences between the Pluronic-coated and Matrigel-coated glass coverslip conditions (Figure 1B1-B2; Figure S2A-B; Movie S1A). On Pluronic-coated glass, around 50% of isolated cells exhibited self-sustained Her1-Venus oscillations (Figure 1B1; Figure 1C1; Figure 1D1; Figure 1G, red), while on Matrigel-coated glass, a majority of the cells were non-oscillatory (Figure 1B2; Figure 1C2; Figure 1D2), with only about 4% of the cells exhibiting oscillations (Figure 1G, blue).

**Figure 1:**
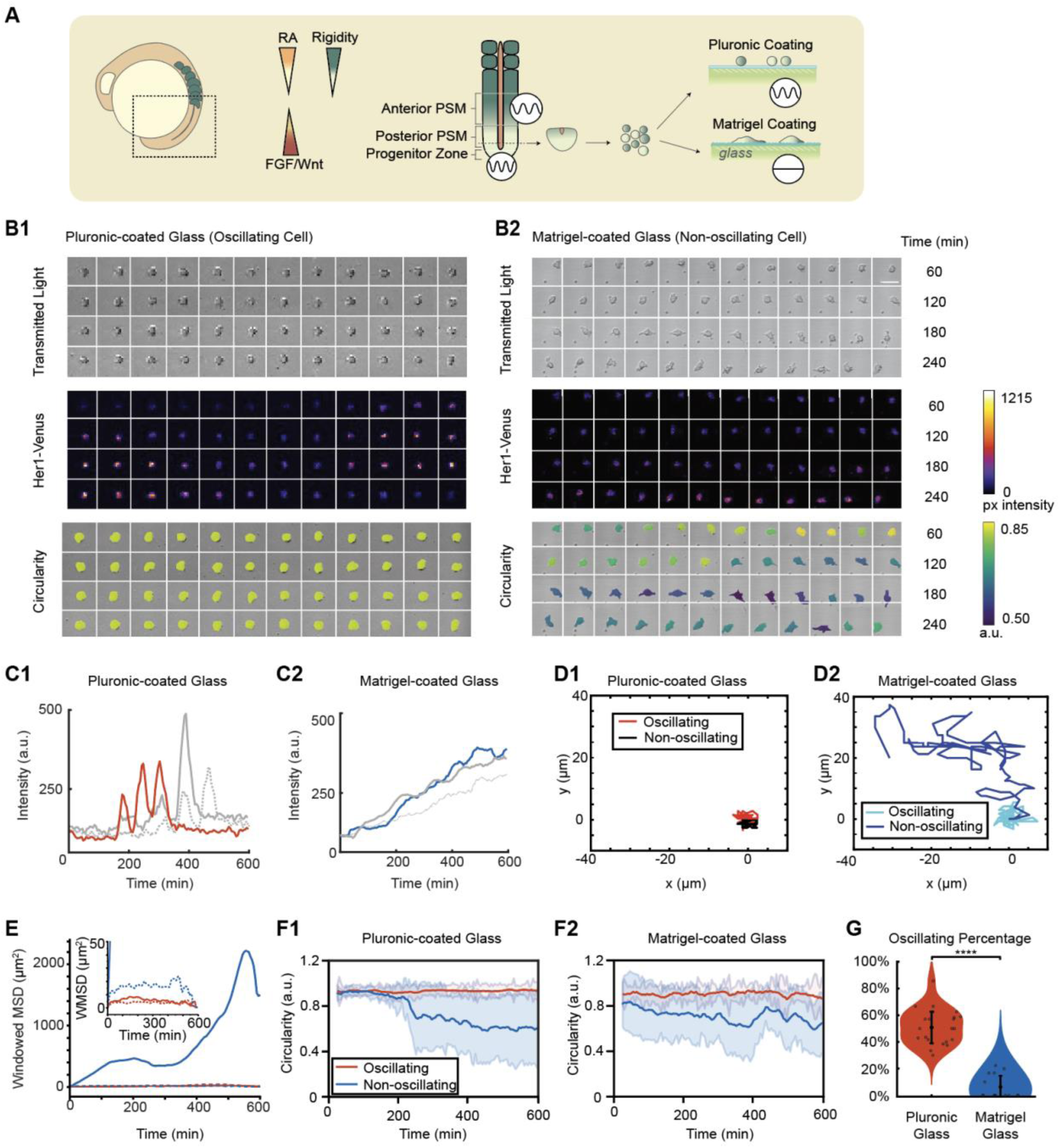
Her1 negative-feedback loop exhibits sustained limit cycle oscillations on low-adhesion surfaces. (A) Schematic of the zebrafish tail during somitogenesis. The segmentation clock in the presomitic mesoderm is known to respond to morphogen gradients (RA, Fgf) and a potential extracellular matrix-mediated mechanical gradient. Progenitor cells harvested from the tailbud are exposed to surfaces with varying rigidities. (B1-B2) Montage of a single cell on Pluronic (B1) and Matrigel-coated glass surfaces (B2). Transmitted light indicates cell viability, Her1-venus intensities indicate oscillations, and circularity demonstrates the PSM cell maintaining a low contact area and spherical conformation on the Pluronic-coated glass surface, while completely spreading on the Matrigel-coated glass surface. Scale bar: 25 µm. (C1-C2) Her1-Venus expression time traces of individual cells on a Pluronic- (hydrophobic) coated glass surface (C1) and Matrigel-coated glass surface (C2). The red line indicates the cell shown in the Pluronic montage panel. Gray lines represent cells from independent experiments. The blue line indicates the cell shown in the Matrigel montage panel. (D1-D2) Tracked cell trajectories of an oscillating and non-oscillating cell on Pluronic- and Matrigel-coated surfaces. Red and blue lines represent cells shown in the respective montage panels. (E) Windowed mean squared displacement (WMSD) of oscillating and non-oscillating cells on Pluronic and Matrigel-coated glass surfaces. The inset provides an enlarged view of the WMSD to highlight the region with low values. Solid red line: oscillating cell on Pluronic-coated glass; Dotted red line: non-oscillating cell on Pluronic-coated glass; Solid blue line: non-oscillating cell on Matrigel-coated glass; Dotted blue line: oscillating cell on Matrigel-coated glass. Solid red and blue lines indicate the respective cells shown in the montages in (B1-B2). (F1-F2) Circularity traces of isolated oscillating (red) and non-oscillating (blue) cells on Pluronic-coated and Matrigel-coated glass surfaces. The plots represent moving average ± SD (the span). The number of tracked cells (n) for circularity in each condition are as follows: Pluronic-coated glass oscillating cells: n= 16. Pluronic-coated glass non-oscillating cells: n= 7. Matrigel-coated glass oscillating cells: n= 7. Matrigel-coated glass non-oscillating cells: n= 9. (G) The percentage of oscillating cells on Pluronic and Matrigel-coated surfaces across experiments. Statistical significance was assessed using two-sample t-tests (two-tailed) comparing oscillation percentages between conditions. The total number of cells analyzed in the two conditions was n = 249 and n = 151, respectively. *P < 0.05, **P < 0.01, ***P < 0.001, ****P < 0.0001.

Moreover, on Pluronic-coated glass, both oscillating (Figure 1D1, red; Figure 1E, solid red) and non-oscillating (Figure 1D1, black; Figure 1E, dotted red) cells exhibited relatively low mean squared displacement (MSD), consistent with Pluronic-coated glass surfaces being not adhesive and not supportive for cell migration. However, on Matrigel-coated glass, non-oscillating cells demonstrated significantly greater cell migration areas (Figure 1D2, blue) and MSD values (Figure 1E, solid blue) that were orders of magnitude higher compared to the oscillating cells (Figure 1D2, light blue; Figure 1E, dotted blue) on the same surface. In both Pluronic-coated and Matrigel-coated conditions, oscillating cells maintained high circularity over time (Figure 1F1-F2, red), while non-oscillating cells eventually became elongated after cell seeding (Figure 1F1-F2, blue), suggesting a potential relation between cell shape, polarity, and oscillatory state. Although Pluronic-coated surfaces are designed to prevent adhesion, non-oscillating cells initially remained rounded but gradually overcame this non-adhesive environment (Figure 1F1, blue), likely through secretion of extracellular matrix proteins, ultimately enabling adhesion, spreading, and migration comparable to non-oscillating cells on Matrigel-coated substrates. These findings suggest that the surface conditions significantly influence the oscillatory behavior of isolated P-PSM cells. Despite their autonomous oscillation capacity, P-PSM cells could lose oscillatory behavior when attached to Matrigel-coated glasses (Figure 1G; Figure S1B). Overall, the observed differences between low-adhesion Pluronic-coated and high-adhesion Matrigel-coated glass conditions suggest that mechanical cues associated with cell adhesion and spreading play a potential role in regulating the segmentation clock. Similar substrate-dependent flattening and loss of oscillations have been reported in chemically dissociated zebrafish^26^ and mouse PSM cells^11^, suggesting that this response is likely a conserved biological phenomenon across species rather than an assay-specific artifact. However, these experiments were performed on ultra-rigid glass coverslips (elastic modulus of 65 GPa^27^), while reported tissue stiffness values across the progenitor zone (PZ), P-PSM, and A-PSM range from 0.3 to 0.8 kPa, as measured using ferrofluid droplet-based approaches in live zebrafish embryos^12,13^. These values are consistent with our atomic force microscopy (AFM) measurements obtained from dissected PZ, P-PSM, and A-PSM tissue explants (Figure S3). Notably, dissected somite tissue exhibited significantly higher stiffness of 2.98 ± 1.21 kPa (Figure S3).

To investigate how the Her1-Venus oscillatory behavior may change across a gradient of rigidity covering the physiologically relevant range, we cultured P-PSM cells on Matrigel-coated PDMS micropost arrays with varying stiffness: 0.6 kPa, 2.9 kPa, 6 kPa, and 1.2 MPa^24^ (Figure 2A; Movie S2A) as well as on Pluronic-coated and Matrigel-coated glass coverslips. P-PSM cells on the two softest PDMS micropost surfaces (0.6 and 2.9 kPa) maintained a high percentage of oscillations, about 40-50%, comparable to those on the Pluronic-coated glass coverslips. However, as the rigidity of the PDMS microposts increased from 2.9 to 6 kPa, the percentage of oscillating P-PSM cells sharply declined to approximately 20%, and this value remained low at higher rigidities (Figure 2B; Figure S4A-B). This suggests a critical rigidity threshold on the order of a few kPa, where PSM cells are most sensitive to mechanical variations in their microenvironment, determining whether they oscillate or not. The cycle number of oscillating P-PSM cells exhibited substantial variability on the softest PDMS microposts (0.6 and 2.9 kPa), with some cells maintaining self-sustained limit-cycle oscillations for up to 17 cycles (Movie S3), whereas cells that managed to oscillate on stiffer PDMS microposts (6 kPa and 1.2 MPa) showed cycle-number distributions that were not significantly different from those observed on Pluronic-coated glass (Figure 2C).

**Figure 2:**
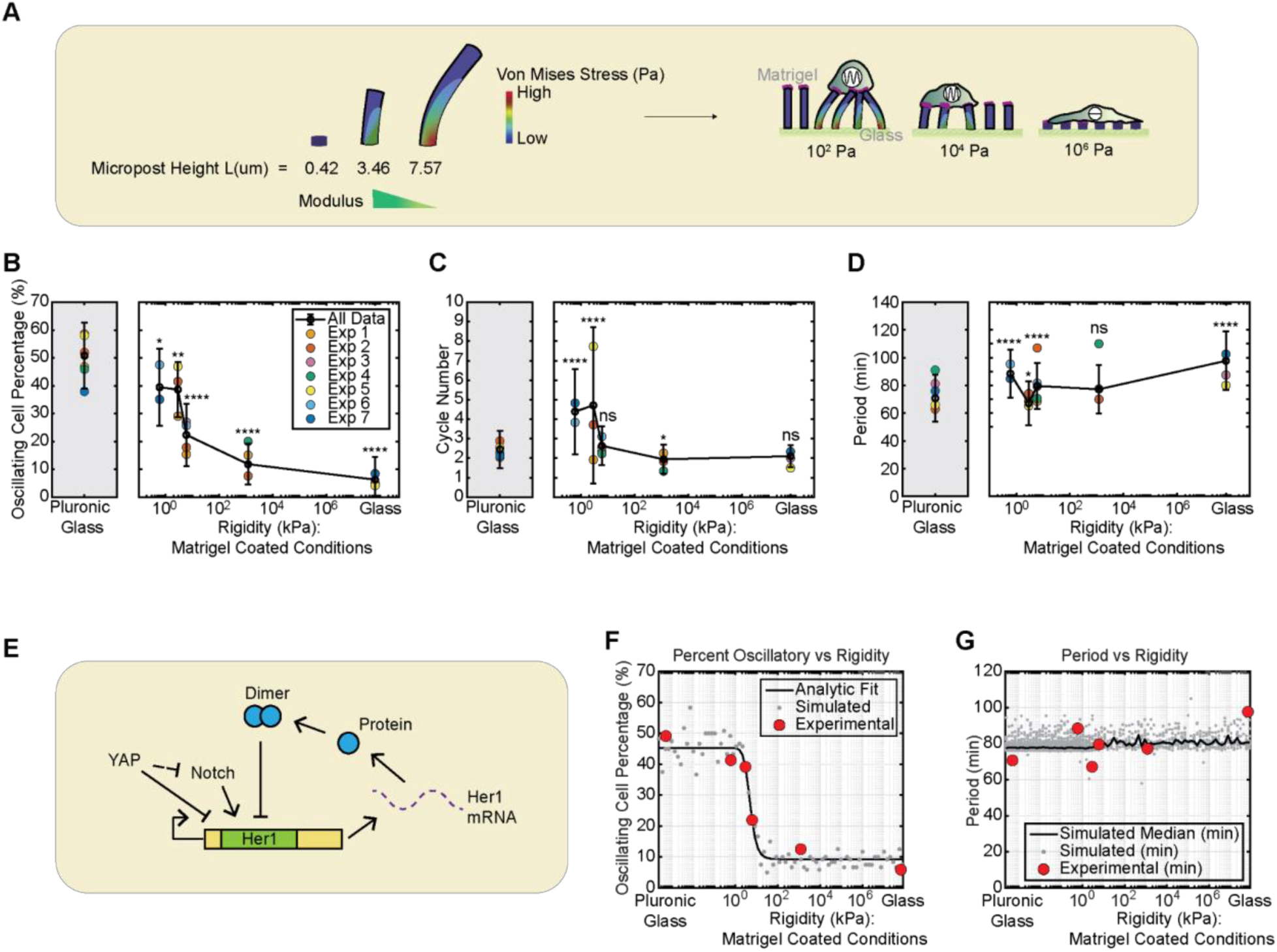
Increasing substrate rigidity reduces the percentage of oscillating cells without a clear effect on the segmentation clock period in zebrafish P-PSM cells isolated using mechanical dissociation. (A) Schematic of the PDMS micropost arrays. Single cells from zebrafish tailbuds were cultured on micropost arrays with varying Young’s modulus: 0.6 kPa, 2.9 kPa, 6 kPa, 1.2 MPa. Pluronic and Matrigel coatings on glass served as extreme controls for the rigidity range, with Matrigel-coated glass exhibiting the highest rigidity and Pluronic-coated glass displaying low cell adhesion. (B-D) Semi-log plots showing the percentage of oscillating cells (%) (B), oscillation cycle number (C), and segmentation clock period (D) in P-PSM cells dissociated across varying substrate rigidities. Grey and white backgrounds indicate Pluronic vs Matrigel surface coatings, respectively. Statistical significance between rigidity conditions was evaluated using pairwise two-sample Welch’s t-tests (two-tailed) assuming unequal variances (ns, not significant; *P < 0.05, **P < 0.01, ***P < 0.001, ****P < 0.0001). For clarity, only selected comparisons between each Matrigel-coated rigidity condition and the Pluronic control are shown; complete pairwise comparisons and exact P-values are provided in the matrix table in Supplementary Table S2. In all panels, black open circles with error bars represent the mean ± SD of the pooled dataset. Colored symbols indicate results from individual independent experiments and follow the color-blind-friendly Okabe-Ito palette^43^. Because experimental constraints limited each experiment to a maximum of four conditions measured simultaneously, data were pooled across N = 7 independent experiments (with at least two replicates per stiffness condition). The number of positions and tracked cells (n) pooled for each experiment is as follows: Exp1: 1.2MPa: total 3 positions, pooled total cells n=53; for each position n1=26, n2=18, n3=9. 6kPa: total 2 positions, pooled total cells n=52; for each position n1=17, n2=35. 2.9kPa: total 4 positions, pooled total cells n=86; for each position n1=10, n2=18, n3=28, n4=30. Pluronic glass: total 3 positions, pooled total cells n=47; for each position n1=23, n2=10, n3=14. Exp2: 1.2MPa: total 4 positions, pooled total cells n=79; for each position n1=4, n2=26, n3=25, n4=24. 6kPa: total 2 positions, pooled total cells n=28; for each position n1=14, n2=14. 2.9kPa: total 6 positions, pooled total cells n=53; for each position n1=11, n2=6, n3=10, n4=7, n5=11, n6=8. Pluronic glass: total 4 positions, pooled total cells n=52; for each position n1=14, n2=9, n3=13, n4=16. Exp3: Matrigel glass: total 4 positions, pooled total cells n=40; for each position n1=12, n2=9, n3=10, n4=9. 6kPa: total 3 positions, pooled total cells n=52; for each position n1=11, n2=19, n3=22. Pluronic glass: total 4 positions, pooled total cells n=29; for each position n1=3, n2=12, n3=7, n4=7. Exp4: 1.2MPa: total 4 positions, pooled total cells n=20; for each position n1=7, n2=4, n3=3, n4=6. 6kPa: total 2 positions, pooled total cells n=35; for each position n1=7, n2=28. Pluronic glass: total 5 positions, pooled total cells n=37; for each position n1=4, n2=6, n3=7, n4=8, n5=12. Exp5: Matrigel glass: total 5 positions, pooled total cells n=48; for each position n1=4, n2=9, n3=12, n4=11. n5=12. 2.9kPa: total 8 positions, pooled total cells n=81; for each position n1=11, n2=9, n3=1, n4=13, n5=10, n6=14, n7=11, n8=12. Pluronic glass: total 5 positions, pooled total cells n=62; for each position n1=17, n2=8, n3=12, n4=16, n5=9. Exp6: 6kPa: total 5 positions, pooled total cells n=79; for each position n1=21, n2=23, n3=15, n4=11, n5=9. 0.6kPa: total 7 positions, pooled total cells n=40; for each position n1=8, n2=1, n3=9, n4=4, n5=6, n6=2, n7=10. Exp7: Matrigel glass: total 7 positions, pooled total cells n=71; for each position n1=6, n2=4, n3=9, n4=10, n5=10, n6=21, n7=11. 0.6kPa: total 6 positions, pooled total cells n=74; for each position n1=9, n2=6, n3=24, n4=15, n5=8, n6=12. Pluronic glass: total 3 positions, pooled total cells n=29; for each position n1=9, n2=10, n3=10. (E) Illustration for a proposed YAP-mediated regulatory mechanism acting on the Her1 negative feedback loop in the segmentation clock. (F) Time-delayed genetic oscillator model fitting data showing the predicted percentage of oscillating cells (%) versus substrate rigidity (kPa). Red dots: experimental data from (B). Black line: analytic model fit. Grey dots: oscillation fractions obtained from Monte-Carlo simulations. (G) Comparison of model fit parameters predicting the clock period (min) versus rigidity (kPa). Red dots: experimental data from (D). Grey dots: individual simulated sample periods. Black line: median simulated period. All fitting parameter values are reported in Supplementary Table S4.

We next examined the distribution of oscillation periods under various substrate rigidity conditions. Dissociated P-PSM cells on Pluronic-coated glass coverslips exhibited a wide distribution of oscillation periods (70.8 ± 17.0 minutes, mean ± SD), longer and more variable compared to in vivo segmentation clock oscillations, but consistent with the reported values from chemically dissociated P-PSM cells cultured in vitro^4^. P-PSM cells on different PDMS micropost arrays exhibited a broad range of oscillation periods that are comparable to those on Pluronic-coated glass coverslips, and these oscillation periods did not exhibit a clear dependency on substrate rigidity (Figure 2D). These data suggest that while cells can transition between quiescent and oscillatory states in response to mechanical cues, the period of the segmentation clock may depend on an intrinsic timing mechanism that is robust to mechanical perturbations. The pie charts in Figure S5 provide a comprehensive overview of the distribution of cell behaviors across different experimental conditions in this study.

To explain the switch-like decrease in the percentage of oscillating P-PSM cells with increasing substrate rigidity, we developed a minimal phenomenological model that links rigidity-dependent YAP activity to the *her1* delayed negative-feedback oscillator (Figure 2E). This model was motivated by a previous study that described the mouse segmentation clock as an activator-repressor oscillator based on the FitzHugh-Nagumo (FHN) model and proposed that the Yap pathway modulates the excitability threshold, effectively acting as a gate for Notch signaling as an external current^11^. We modeled zebrafish single-cell Her1 dynamics using a time-delayed genetic oscillator model^28^, assuming Her1 protein half-life of 5 min and a negative feedback delay of 35 min (including transcription and translation). These values were chosen to recapitulate the slower oscillatory dynamics observed in isolated PSM cells (60-100 min), relative to the embryonic clock (30 min), while remaining within literature-reported ranges for Her1 turnover (3.5-7 min) and total feedback delay (13-35 min)^29–32^. Substrate rigidity modulates YAP activity through a Hill function, and YAP (inhibitor) and Notch (activator) antagonistically regulate the Her1 production rate, *χ*. To capture cell-to-cell variability, *χ* was sampled from a Gaussian distribution at each stiffness, and individual simulated trajectories were used to detect oscillations and extract periods (Figure S6A). Fitting this model to the experimentally measured percentage of oscillating cells across substrates yielded a rigidity threshold of 3.77 kPa (95% CI: 2.58-5.37 kPa) (Figure 2F), which falls within the experimentally observed transition range between 2.9 and 6 kPa (Figure 2B) and is also close to the reported switch-like YAP nuclear translocation threshold of 5 kPa in cultured mouse embryonic fibroblasts^20^ (Figure S6B-C). Using the fitted parameters, the model reproduced the sigmoidal response of oscillating-cell percentage to rigidity, while predicting little to no effect of rigidity on the oscillation period (Figure 2G), consistent with experimental observations.

Consistent with this interpretation, YAP1 immunofluorescence showed a significantly higher nuclear-to-cytoplasmic YAP1 ratio in dissociated P-PSM cells cultured on Matrigel-coated glass coverslips compared with those cultured on Pluronic-coated glass coverslips (Figure S7). In addition, under Matrigel-coated glass conditions, YAP inhibition by Verteporfin increased the fraction of oscillating cells relative to DMSO-treated controls, with little consistent effect on oscillation period (Figure S8). Together, these results support a role for YAP-associated mechanotransduction in regulating the oscillatory state of PSM cells, while suggesting that YAP activity has a limited influence on the intrinsic period of cells that remain oscillatory.

### Oscillating and Non-Oscillating PSM Cells Exhibit Distinct Biophysical Properties

We further explored the relationship between morphology and oscillatory behavior of P-PSM cells on substrates of varying rigidities. Similar to our observations from Pluronic-coated and Matrigel-coated glass coverslips, oscillating and non-oscillating P-PSM cells exhibited distinct morphological and biophysical properties on PDMS micropost arrays of varying rigidities. Across all tested rigidities (0.6 kPa - 1.2 MPa), and consistent with glass coverslip conditions (Figure 1F1-F2), oscillating P-PSM cells consistently exhibited greater and more persistent circularity than their non-oscillating counterparts (Figure 3A1-A4; Figure S9; Figure S10A-B). Distinct changes in cell circularity were associated with oscillatory behaviors of P-PSM cells, with actively oscillating cells maintaining high circularity (Figure 3B1) and non-oscillating cells showing a significant reduction in circularity (Figure 3B2). Additionally, a greater percentage of P-PSM cells showed a rounded morphology on Matrigel-coated glass coverslips than on Matrigel-coated 1.2 MPa micropost arrays (Figure S10A-B), indicating that the morphology of isolated P-PSM cells was likely sensitive to surface topography. Together, PSM cells took longer to spread on softer substrates, while non-oscillating cells exhibited a more rapid decrease in circularity as substrate rigidity increased (Figure 3C).

**Figure 3:**
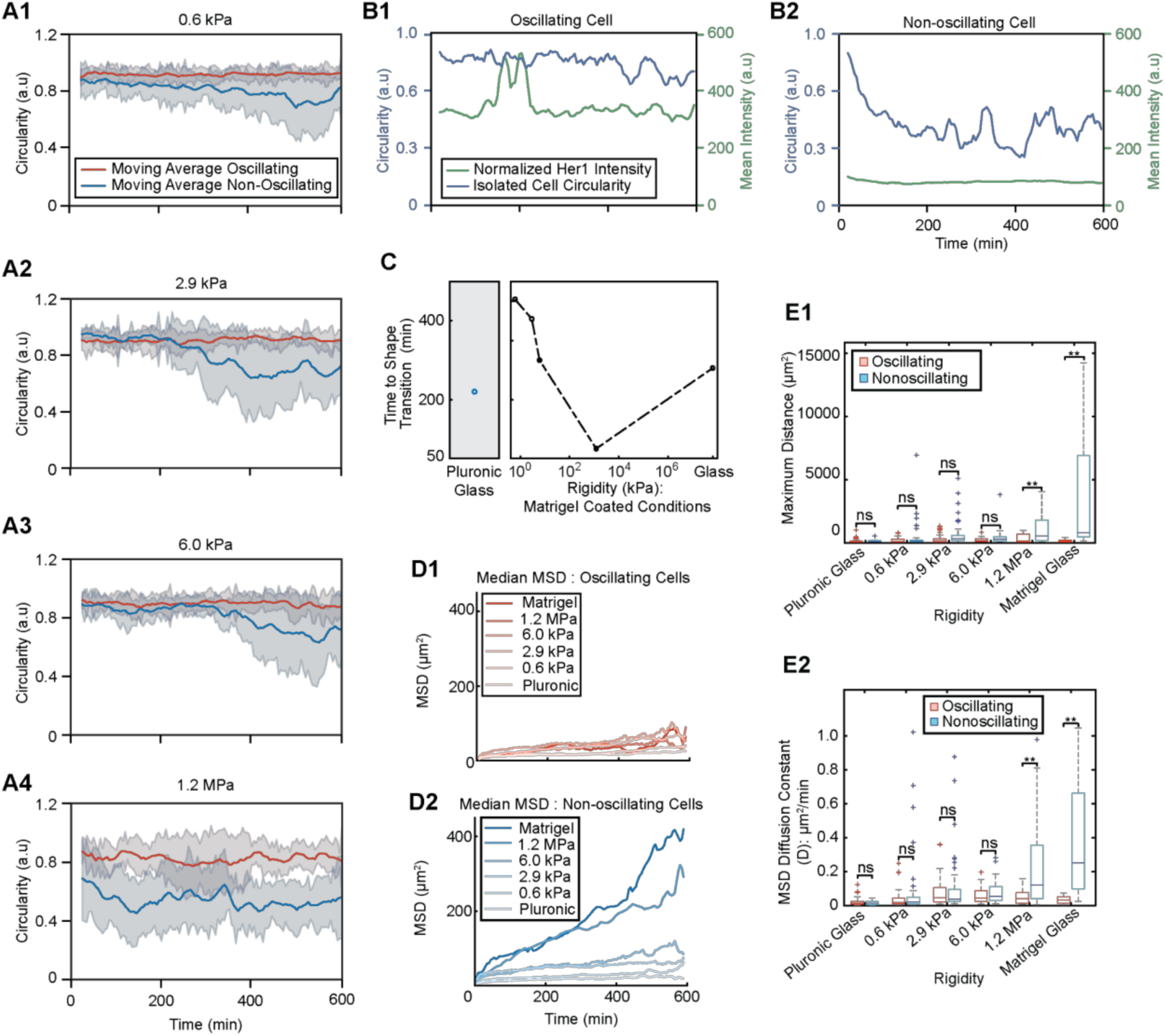
Substrate rigidity modulates morphological dynamics and migratory behavior in isolated PSM cells exhibiting distinct oscillation states. (A1-A4) Single-cell circularity traces over time on 0.6 kPa (A1), 2.9 kPa (A2), 6 kPa (A3), and 1.2 MPa (A4) surfaces, for oscillating cells (red) and non-oscillating cells (blue). The plots represent moving average ± SD (the span). The number of tracked oscillating/non-oscillating cells (n) for each condition in A1-A4 are as follows: 0.6kPa (n=10/10), 2.9kPa (n=10/7), 6.0kPa (n=10/7), 1.2MPa (n=6/7). (B1-B2) Circularity and Her1-Venus intensity traces of single cells on a 1.2 MPa micropost-array surface. The oscillating cell (n=1) (B1) maintains high circularity, while the non-oscillating cell (n=1) (B2) progressively loses circularity. (C) Time for single non-oscillating cells to reach a distinct geometry (change in circularity) from their oscillating counterpart. (D1-D2) Mean squared displacement (MSD) analyses of cells on varying surface rigidities, including median MSD of oscillating (D1) and non-oscillating (D2) cells. (E1) Maximum squared distance traveled by oscillating cells and non-oscillating cells, where the plots represent the mean ± SD. (E2) MSD-derived diffusion coefficient D of oscillating and non-oscillating cells among different rigidity, where the plots represent the mean ± SD. Oscillating/non-oscillating cells (n) for each condition in D1-2 and E1-2: Pluronic (n=43/22), 0.6kPa (n=38/40), 2.9kPa (n=58/88), 6.0kPa (n=28/43), 1.2MPa (n=10/48), Matrigel (n=5/25). Statistical significance is assessed using a Wilcoxon rank-sum test (two-tailed) based on the total counts for each condition. *P < 0.05, **P < 0.01, ***P < 0.001. Complete P-value matrix table is in Table S1: Supplementary Data Table for Figure 3.

Analysis of MSD revealed that oscillating P-PSM cells consistently maintained low MSD values across all substrate rigidity conditions, while non-oscillating ones showed a significant increase in MSD over time, with a steeper rise on stiffer surfaces (Figure S10C-D; Median MSD: Figure 3D1-D2; Maximum squared distance: Figure 3E1). These differences are further demonstrated in the MSD diffusion coefficient (D) analysis, which shows that oscillating P-PSM cells consistently maintain low D values regardless of substrate rigidity conditions. In contrast, non-oscillating P-PSM cells exhibit increased D with higher substrate rigidity values (Figure 3E2).

To provide a detailed view of cell-substrate interactions for oscillating and non-oscillating P-PSM cells, we captured higher-resolution images of P-PSM cells on 0.6 kPa and 2.4 kPa micropost arrays and analyzed the traction forces they exerted on the substrates as they attached and migrated. The subcellular-level traction forces were quantified based on the deflections of the micropost tops^24^. Oscillating P-PSM cells on 0.6 kPa micropost arrays maintained a round shape and exerted low traction forces (Figure 4A; Figure 4C, red; Movie S4A), whereas non-oscillating P-PSM cells on the same substrate became polarized and increased traction forces approximately 4 hours after seeding (Figure 4B; Figure 4C, blue; Movie S4B). These differences were also observed for oscillating and non-oscillating PSM cells on the 2.4 kPa micropost arrays (Movies S4C-D). The traction force normalized by cell spread area indicated that oscillating cells exhibited lower contractility compared to non-oscillating cells on both 0.6 kPa and 2.4 kPa micropost arrays (Figure 4D-E).

**Figure 4:**
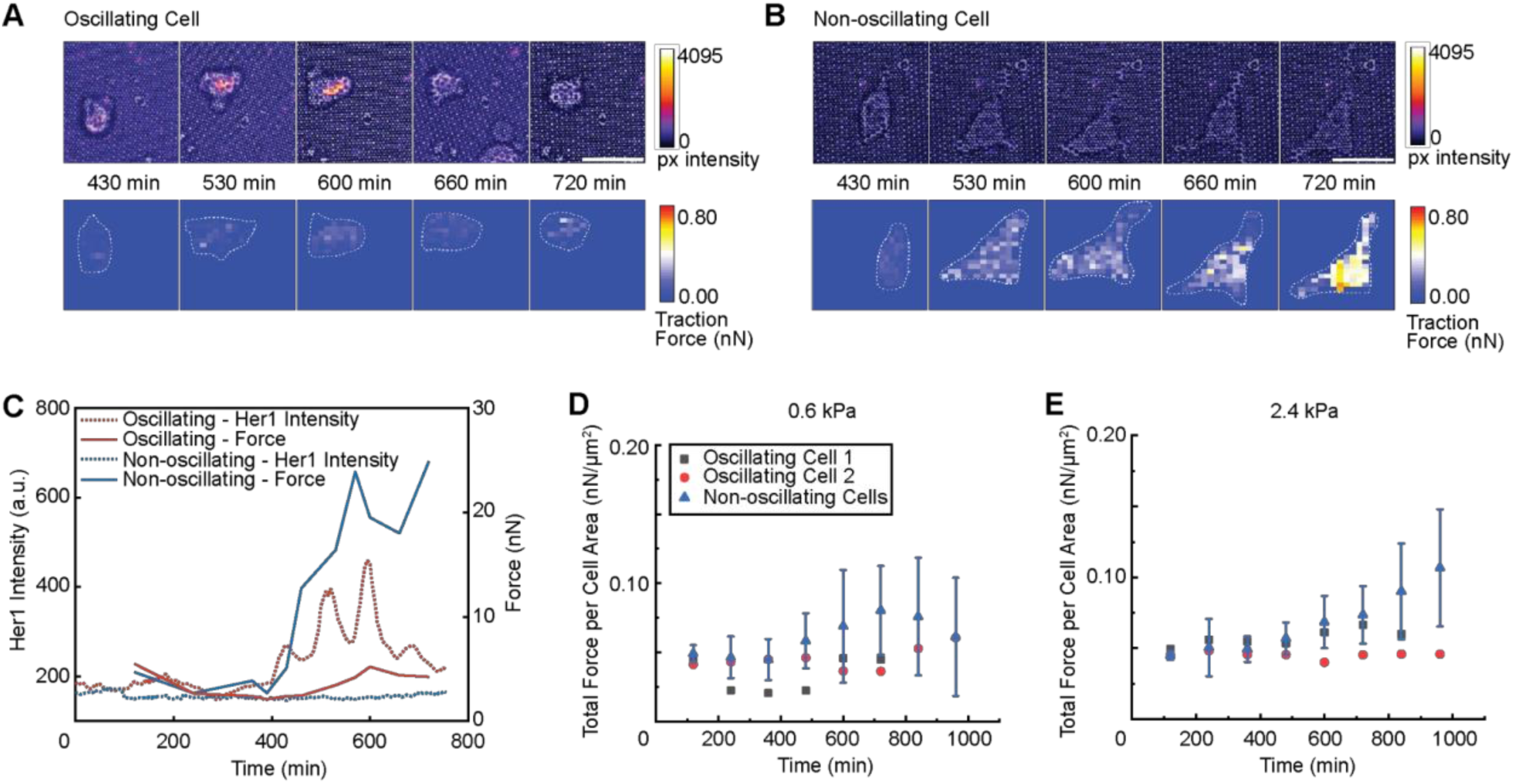
Oscillating and non-oscillating PSM cells exhibit distinct traction force profiles across substrate rigidities. (A-B) Her1-Venus intensity and traction force heat maps of oscillating (A) and non-oscillating (B) isolated cells on 0.6 kPa micropost arrays. (C) Her1 intensity and force profiles over time for the oscillating (red) and non-oscillating (blue) cells. (D) Quantitative analysis of total force per cell area for oscillating (cell number n = 2) and non-oscillating (n = 5) isolated PSM cells cultured on 0.6 kPa micropost arrays, where the plots represent the mean ± SD. (E) Quantitative analysis of total force per cell area for oscillating (n = 2) and non-oscillating (n = 5) isolated PSM cells cultured on 2.4 kPa micropost arrays, where the plots represent the mean ± SD. Scale bar: 15 µm.

These findings indicate that cell morphology, spreading dynamics, motility, and mechanical tension are correlated with oscillatory behaviors of the segmentation clock. The observed morphological and biophysical differences between oscillating and non-oscillating PSM cells may be associated with YAP activation, which has been reported to respond to mechanical cues and cell stretching in both mouse cell cultures^11,20^ and zebrafish PSM cells (Figure S7). Collectively, these results suggest that mechanical cues from the microenvironment could potentially influence the oscillatory dynamics of the segmentation clock through changes in cell shape, contractility, and mechanical tension.

### Multicellular Aggregates Display Emergent Oscillatory Properties Influenced by Cell-Cell Interactions and Mechanical Cues

To investigate the oscillatory behavior of PSM cells in a multicellular context, we cultured aggregates of P-PSM cells on PDMS micropost arrays and compared their properties to those of single cells across different substrate rigidity conditions (Figure 5A). Similar to isolated P-PSM cells, aggregates of P-PSM cells displayed oscillatory behaviors linked to morphology and substrate conditions, suggesting the influence of mechanical signals in cell clusters. As an example, clusters of P-PSM cells could either oscillate (Movie S5A) or not oscillate (Movie S5B) on Pluronic-coated glass coverslips, 2.9 kPa micropost arrays, and 1.2 MPa micropost arrays. In some clusters on 1.2 MPa micropost arrays, cells that initially oscillated at the center migrated to the periphery, where they lost oscillations and displayed increased cellular spreading (Movie S5C). On Pluronic-coated glass coverslips, cell colonies were able to maintain sustained Her1-Venus oscillations along with stable circularity and morphology over time, without spreading (Figure 5B-C, red; Movie S5A, Pluronic glass). There was a slight reduction in circularity, primarily due to the protrusion of some peripheral cells (Figure 5D1). Similar behaviors were observed on soft substrates (Movie S5A, 2.9 kPa). Conversely, on 1.2 MPa PDMS microposts, cell aggregates were more prone to stretching and were predominantly non-oscillatory, except for transient oscillations in some central cells before they migrated and spread. These aggregates experienced a significant elongation and reduction in circularity over time (Figure 5B-C, blue; Figure 5D2; Movie S5C).

**Figure 5:**
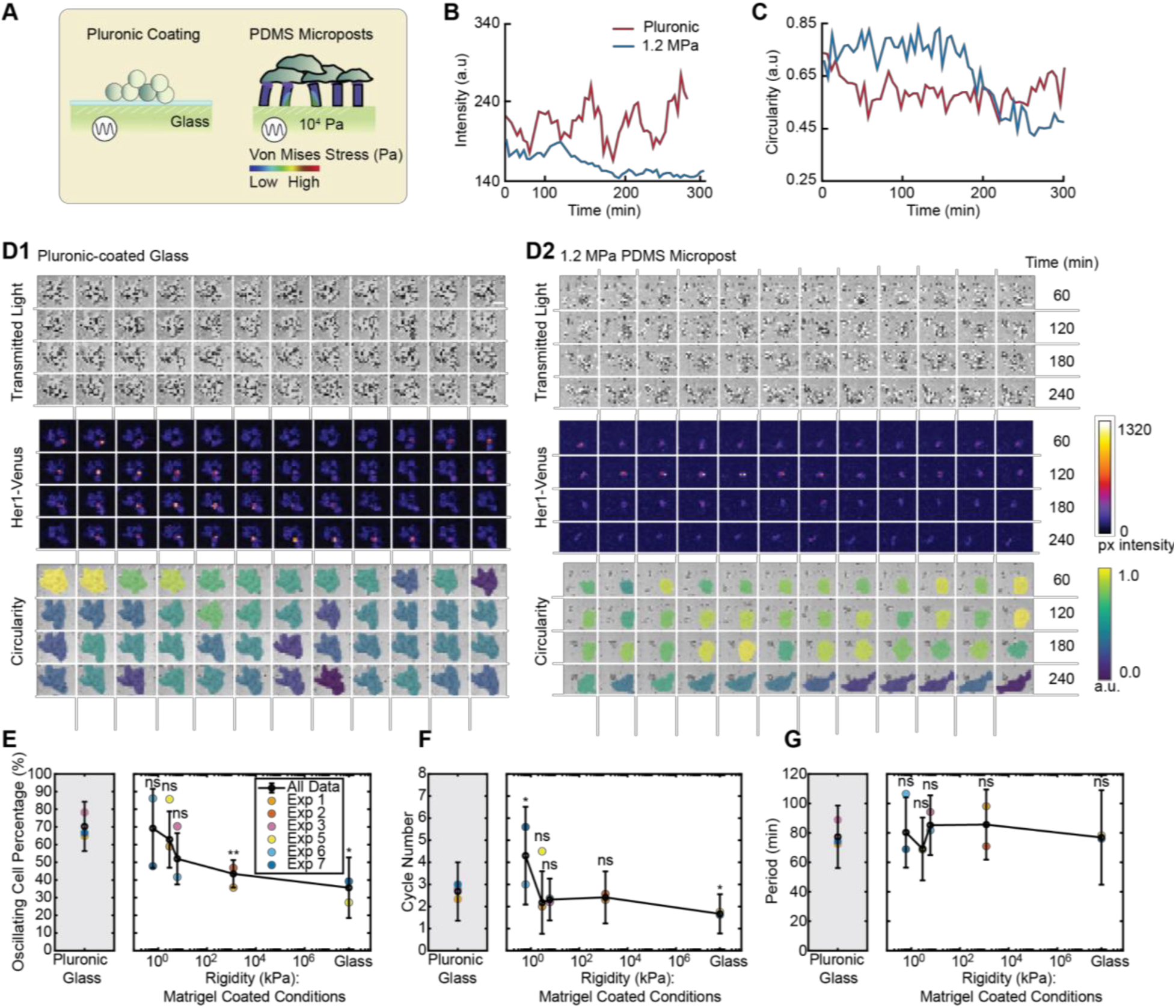
Oscillatory behavior and morphological properties of PSM cell aggregates across rigidity conditions. (A) Schematic of cell aggregates cultured on PDMS micropost arrays. (B) Her1-Venus intensity traces of cell aggregates on Pluronic-coated glass surfaces (black) and 1.2 MPa PDMS micropost arrays (red). (C) Circularity traces of the representative cells in (B). (D1) Montage of an oscillating cell aggregate on Pluronic-coated glass. (D2) Montage of a heterogeneous aggregate on 1.2MPa micropost arrays, which segregated into two layers, with oscillatory cells in the upper layer and elongated, non-oscillatory cells in the lower layer (Movie S5C). Transmitted light indicates aggregate viability, Her1-Venus intensities reflect segmentation clock oscillations, and circularity demonstrates cell morphology. Scale bar: 40 µm. (**E**) Semi-log plot of the percentage of oscillating cell aggregates on surfaces of varying rigidities. (**F**) Semi-log plot showing oscillating cell cycle number of aggregates versus surface rigidity (kPa). Grey and white background shading indicate Pluronic-coated and Matrigel-coated surfaces, respectively. (**G**) Semi-log plot showing segmentation clock period (min) of aggregates versus surface rigidity (kPa). Grey and white background shading indicate Pluronic-coated and Matrigel-coated surfaces, respectively. Statistical significance between rigidity conditions was evaluated using pairwise two-sample Welch’s t-tests (two-tailed) assuming unequal variances (ns, not significant; *P < 0.05, **P < 0.01, ***P < 0.001, ****P < 0.0001). The plots of (E-G) represent the mean ± SD. The number of positions and tracked aggregates (n) pooled for each experiment is as follows: Exp1: 1.2MPa: total 3 positions, pooled total aggregates n=28; for each position n1=13, n2=8, n3=7. 2.9kPa: total 4 positions, pooled total aggregates n=83; for each position n1=13, n2=17, n3=20, n4=33. Pluronic glass: total 3 positions, pooled total aggregates n=20; for each position n1=8, n2=5, n3=7. Exp2: 1.2MPa: total 3 positions, pooled total aggregates n=64; for each position n1=9, n2=30, n3=25. Exp3: 6kPa: total 1 position, pooled total aggregates n=27. Pluronic glass: total 1 position, pooled total aggregates n=23. Exp5: Matrigel glass: total 2 positions, pooled total aggregates n=22; for each position n1=11, n2=11. 2.9kPa: total 1 position, pooled total aggregates n=14. Exp6: 6kPa: total 3 positions, pooled total aggregates n=48; for each position n1=20, n2=20, n3=8. 0.6kPa: total 2 positions, pooled total aggregates n=29; for each position n1=7, n2=22. Exp7: Matrigel glass: total 6 positions, pooled total aggregates n=51; for each position n1=9, n2=7, n3=10, n4=2, n5=11, n6=12. 0.6kPa: total 3 positions, pooled total aggregates n=23; for each position n1=9, n2=9, n3=5. Pluronic glass: total 1 position, pooled total aggregates n=21.

We analyzed the oscillation properties of cell aggregates across all substrate rigidity conditions. Compared with single cells, aggregates exhibited a generally higher percentage of oscillating cells across all conditions, with substantially reduced sensitivity to substrate rigidity (Figure 5E). The switch-like decrease in oscillating cell percentage observed in single cells was largely absent in aggregates: even on stiff substrates (1.2 MPa and Matrigel-coated glass), the reduction in oscillating fraction was modest. In addition, neither oscillation cycle number nor oscillation period displayed a clear dependence on substrate rigidity (Figures 5F and 5G). Aggregates cultured on Pluronic-coated glass exhibited a broad distribution of oscillation periods (77.3 ± 21.2 min, mean ± SD), and this broad distribution was maintained across rigidity conditions, overlapping with the 60-100 min range observed in single cells and consistent with an intrinsic timing mechanism (Figure 5G). Together, these results suggest that while single-cell oscillatory competence is strongly gated by substrate mechanics, cell-cell communication within aggregates buffers this mechanical sensitivity and broadly promotes the oscillatory state, pointing to a role for Notch - mediated intercellular signaling in maintaining clock activity against the local mechanical constraints.

These findings highlight the complex interplay between cellular interactions and the mechanical environment in regulating the segmentation clock dynamics in multicellular contexts. The observed differences in the oscillatory response to rigidity between aggregates of PSM cells and single PSM cells may be attributed to the enhanced Notch signaling in the aggregates, which positively regulates *her1*, coupled with the antagonistic effect of mechano-transduced YAP activity on the segmentation clock. Another possibility could be the difficulty in activating YAP signaling in cell aggregates compared to isolated cells, due to the inhibited cell spreading or stretching of the cells in the middle. This may also explain the heterogeneous pattern of oscillations observed in aggregates on rigid substrates, where cells at the center are more likely to oscillate while cells in the periphery tend to spread and do not oscillate (Movie S5C).

## Discussion

Our study reveals that the segmentation clock is sensitive to mechanical cues from the microenvironment, with substrate rigidity playing a crucial role in modulating the oscillatory behavior of isolated PSM cells. Notably, we observed a critical rigidity threshold between 2.9 and 6 kPa, where the percentage of oscillating PSM cells exhibits a switch-like drop, suggesting that the segmentation clock is finely tuned to respond to specific tissue mechanical ranges.

The use of PDMS micropost arrays in our study provides a unique and powerful tool to investigate the role of mechanics in regulating the segmentation clock at the single-cell level and in multicellular aggregates. This approach allows for precise control over the mechanical environment, enabling us to explore a wide range of substrate rigidities and their effects on oscillatory behaviors of PSM cells. Hydrogel-based methods to modulate substrate rigidities require adjusting the densities of polymers, which impacts not only rigidity but also ligand concentration and other complex factors like surface topology and matrix porosity that can also affect cell behaviors. These complexities make it difficult to distinguish biochemical effects from mechanical effects on cellular responses. In our study, we cultured cells on micropost arrays with a uniform 2% Matrigel coating across all rigidity conditions to isolate mechanical rigidity from other matrix properties. This approach enables a clearer interpretation of cellular responses specifically to bulk mechanical changes. Additionally, this method could be extended to investigate the role of mechanics in segmentation clock systems across different species, providing a valuable tool for comparative studies and deepening our understanding of the conserved and divergent mechanisms that regulate the segmentation clock across vertebrates.

Our findings also support the possible roles of cell morphology, motility, and mechanical tension in regulating the oscillatory dynamics of the segmentation clock. We observed that oscillating PSM cells exhibit distinct biophysical properties, such as sustained circularity, reduced spreading, lower motility, and decreased mechanical tension. In contrast, non-oscillating PSM cells display altered morphology, increased spreading, higher motility, and elevated traction forces. These results suggest that the mechanical state of individual PSM cells, as well as their ability to sense and respond to mechanical cues from the microenvironment, might be critical factors in determining the oscillatory behavior of the segmentation clock.

Furthermore, our study reveals that cell-cell contacts and the mechanical environment within multicellular aggregates may coordinate, resulting in the segmentation clock of the PSM cell aggregates being less sensitive to rigidity changes compared to isolated PSM cells. These findings underscore the importance of investigating the segmentation clock dynamics in both single-cell and multicellular contexts, as the interplay between cell-cell communication and mechanical cues can give rise to emergent behaviors that are not observed in isolated cells.

All experiments in this study were performed using mechanically dissociated PSM cells to best preserve intrinsic cellular properties, although this approach limited sample size. To increase throughput, we additionally performed chemical dissociation, which yielded comparable results for both P-PSM dissociated cells (Figure S11; Movies S1B and S2B) and cell aggregates (Figure S12; Movie S5D-E), including a consistent switch-like rigidity-dependent reduction in oscillating cell percentage in single cells and its rescue in aggregates.

Our in vitro experiments show oscillation periods ranging from 60 to 100 min in both isolated single cells and small aggregates derived from mechanically dissociated P-PSM cells cultured in serum-free L-15 medium at 28°C. These values are broadly consistent with the dissociated-cell measurements reported by Webb et al.^4^, who observed 62 ± 21 min for fully isolated cells and 78.3 min for low-density dissociated cells in L-15 medium supplemented with 10% FBS and exogenous Fgf8b at 26°C. Remaining differences may arise from variations in temperature (26°C vs. 28°C), medium composition (serum/Fgf8b vs. serum-free), dissociation method (chemical vs. mechanical), and differences in the composition and axial identity of dissociated tailbud cell populations^8^.

While these in vitro assays enable controlled manipulation of mechanical cues by varying substrate rigidity in two-dimensional (2D) environments, they do not recapitulate the full complexity of the in vivo embryonic context, where cells experience three-dimensional (3D) confinement, anisotropic mechanical forces, and tissue-scale deformations. Future studies in more physiologically relevant 3D systems^33–37^ will therefore be required to fully understand how mechanical cues regulate segmentation clock dynamics in vivo.

## Acknowledgment

We thank members of the Yang lab for scientific discussions and feedback on the manuscript and the ULAM staff and vivarium services for fish husbandry. We thank Dr. Sean Megason and members of the lab for sharing knowledge and advice on zebrafish husbandry and embryo imaging. We thank Dr. Sharon Amacher for generously providing the transgenic *Tg(her1:her1-Venus)* zebrafish and Dr. Ertugrul Ozbudak for generously providing the transgenic *Tg(her7:her7-Venus)* zebrafish. We thank Dr. Xufeng Xue for providing training on the fabrication of PDMS micropost arrays and technical support. We thank Dr. Damon Hoff and the Single Molecule Analysis in Real-Time (SMART) Center for technical support with AFM experiments. We thank Danya Rimawi (REU Summer 2025) for assistance in testing a cryosectioning protocol for zebrafish tailbuds, intending to enable AFM measurements across different tailbud regions. This work was supported by grants from the NSF (MCB #2218083), NIH (R21HD105126; R01GM144584), and the Margaret and Herman Sokol Faculty Awards at the University of Michigan.

## Author contributions

QY and JF conceived the project. QY and JF supervised the project. CS and YY planned and executed experiments. CS, YY, UK, OB, LY, and LL performed image analysis and data analysis. DW and QY implemented the mathematical model. UK, CS, and QY wrote the manuscript. CS, YY, UK, OB, LY, QY, JF edited the manuscript.

## Methods

### Reagents and Resources

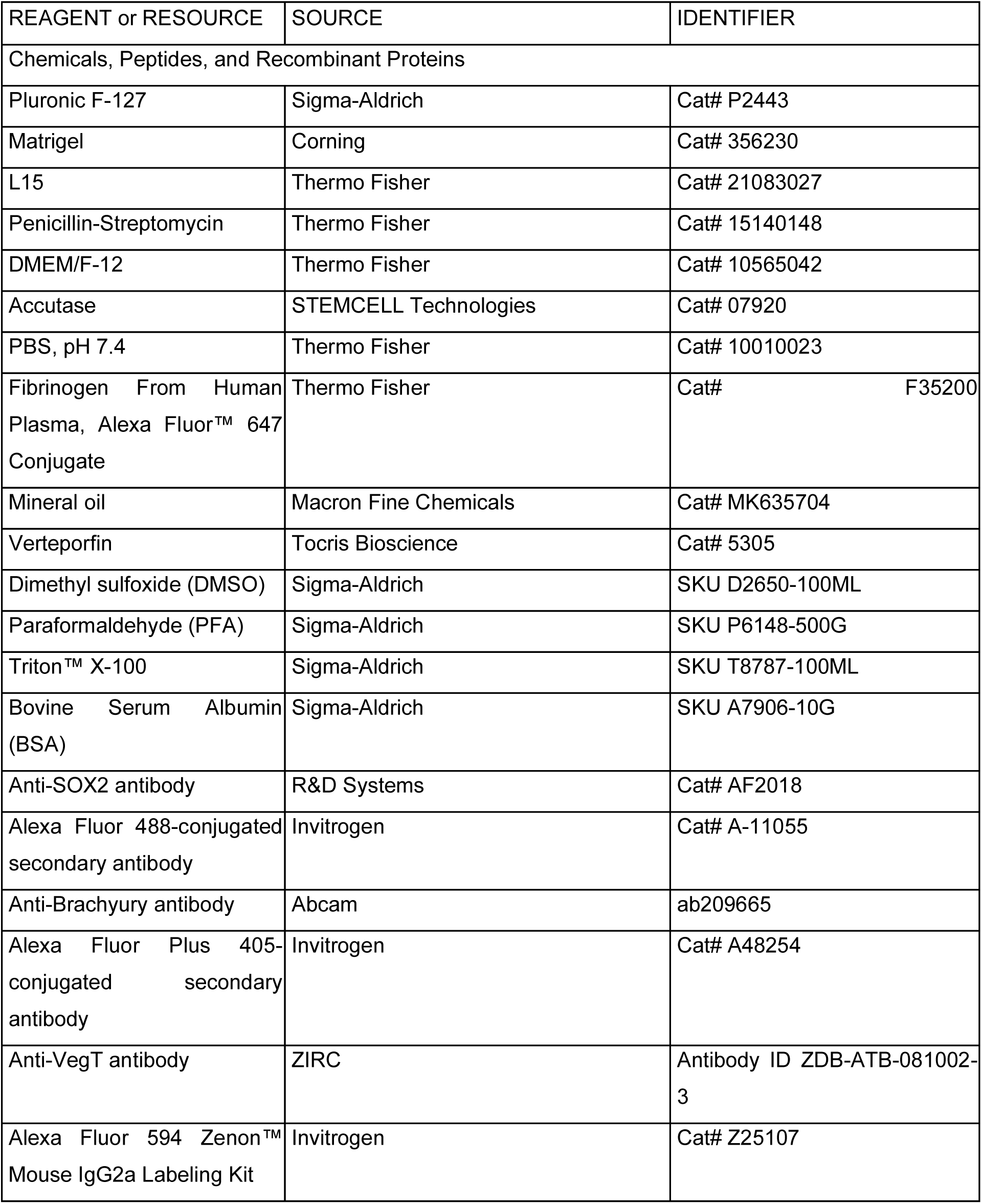
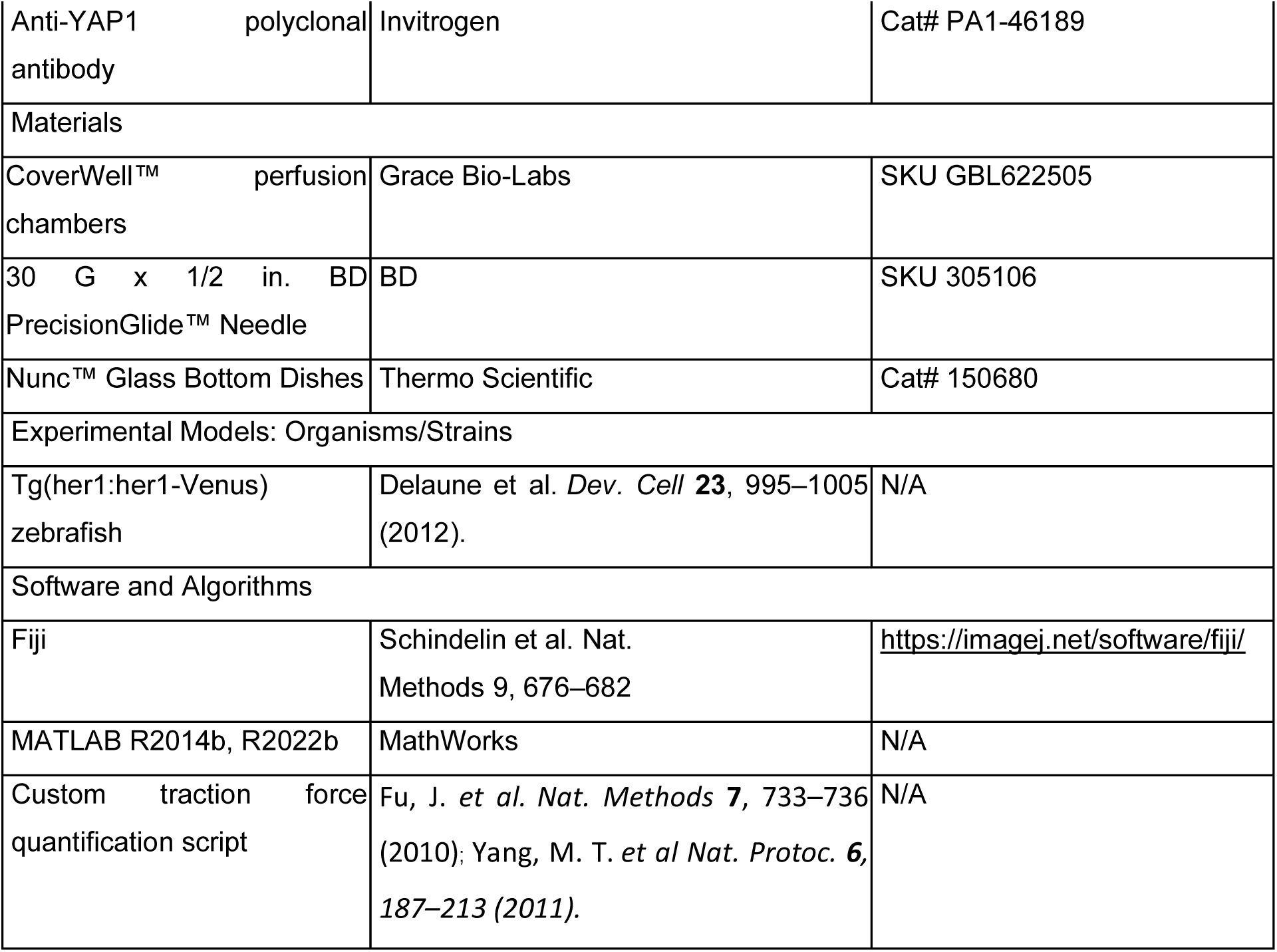

| REAGENT or RESOURCE | SOURCE | IDENTIFIER |
| --- | --- | --- |
| Chemicals, Peptides, and Recombinant Proteins |  |  |
| Pluronic F-127 | Sigma-Aldrich | Cat# P2443 |
| Matrigel | Corning | Cat# 356230 |
| L15 | Thermo Fisher | Cat# 21083027 |
| Penicillin-Streptomycin | Thermo Fisher | Cat# 15140148 |
| DMEM/F-12 | Thermo Fisher | Cat# 10565042 |
| Accutase | STEMCELL Technologies | Cat# 07920 |
| PBS, pH 7.4 | Thermo Fisher | Cat# 10010023 |
| Fibrinogen From Human Plasma, Alexa Fluor™ 647 Conjugate | Thermo Fisher | Cat# F35200 |
| Mineral oil | Macron Fine Chemicals | Cat# MK635704 |
| Verteporfin | Tocris Bioscience | Cat# 5305 |
| Dimethyl sulfoxide (DMSO) | Sigma-Aldrich | SKU D2650-100ML |
| Paraformaldehyde (PFA) | Sigma-Aldrich | SKU P6148-500G |
| Triton™ X-100 | Sigma-Aldrich | SKU T8787-100ML |
| Bovine Serum Albumin (BSA) | Sigma-Aldrich | SKU A7906-10G |
| Anti-SOX2 antibody | R&D Systems | Cat# AF2018 |
| Alexa Fluor 488-conjugated secondary antibody | Invitrogen | Cat# A-11055 |
| Anti-Brachyury antibody | Abcam | ab209665 |
| Alexa Fluor Plus 405-conjugated secondary antibody | Invitrogen | Cat# A48254 |
| Anti-VegT antibody | ZIRC | Antibody ID ZDB-ATB-081002-3 |
| Alexa Fluor 594 Zenon™ Mouse IgG2a Labeling Kit | Invitrogen | Cat# Z25107 |
| Anti-YAP1 polyclonal antibody | Invitrogen | Cat# PA1-46189 |
| Materials |  |  |
| CoverWell™ perfusion chambers | Grace Bio-Labs | SKU GBL622505 |
| 30 G x 1/2 in. BD PrecisionGlide™ Needle | BD | SKU 305106 |
| Nunc™ Glass Bottom Dishes | Thermo Scientific | Cat# 150680 |
| Experimental Models: Organisms/Strains |  |  |
| Tg(her1:her1-Venus) zebrafish | Delaune et al. *Dev. Cell* **23**, 995–1005 (2012). | N/A |
| Software and Algorithms |  |  |
| Fiji | Schindelin et al. *Nat. Methods* **9**, 676–682 | https://imagej.net/software/fiji/ |
| MATLAB R2014b, R2022b | MathWorks | N/A |
| Custom traction force quantification script | Fu, J. *et al. Nat. Methods* **7**, 733–736 (2010); Yang, M. T. *et al Nat. Protoc.* **6**, 187–213 (2011). | N/A |

### Fish husbandry and dissociation for zebrafish tailbud cells

Zebrafish *Tg(her1:her1-Venus)* and *Tg(her7:her7-Venus)* embryos were maintained at 28 °C in E3 buffer without methylene blue until 50% epiboly and subsequently incubated at 19 °C overnight before in vitro experiments. All mechanical dissociation experiments and some chemical dissociation experiments (including exp1 on Pluronic-coated glass, exp1 on 2.9 kPa microposts, exp1 on Matrigel-coated glass) were performed using *Tg(her1:her1-Venus)* embryos. The remaining chemical dissociation experiments utilized *Tg(her7:her7-Venus)*, as this line exhibited improved embryo viability and a stronger fluorescence signal. Embryos were screened at the 5-6 somite stage, and developmental timing was adjusted when needed by changing external temperature to accelerate or decelerate development. In parallel, dissociation reagents and media were prepared: L15 medium supplemented with 0.5% penicillin-streptomycin (PS) (99.5% L15 + 0.5% PS) was warmed to 28 °C, and Accutase was thawed at 28 °C. Matrigel was thawed at 4 - 5 °C, while Pluronic was maintained at room temperature.

For imaging and culturing cells on Pluronic-coated or Matrigel-coated glass, Grace perfusion chambers were assembled using the thinnest available chamber and coverslip. Adhesive was removed, chambers were adhered to coverslips, and edges were pressed to minimize air bubbles. Chambers were washed three times with 70% ethanol, air-dried, and kept clean until use. Each well was coated with either Pluronic or Matrigel, with 40 µL applied per well and incubated for 30 min in a humid environment. Pluronic coating solution was prepared as 0.2% Pluronic (diluted in sterile PBS), and Matrigel coating solution was prepared as 2% Matrigel (diluted in L15). After incubation, coating solutions were removed; Pluronic-coated wells were washed three times with PBS and Matrigel-coated wells were washed three times with L15, with all liquid aspirated after the final wash.

Embryos were dechorionated in E3 buffer manually using two sharp tweezers with minimal shear stress, transferred into L15 medium with penicillin-streptomycin, and yolks were perforated and drained. Tailbuds were dissected on a PDMS-lined dissection dish to minimize mechanical stimuli to embryos, with dissections targeted to collect posterior presomitic mesoderm (P-PSM) and with additional removal of non-target tissues performed as needed. The tissue below the notochord, including the progenitor zone and parts of the posterior PSM, was cut using two syringe needles where one needle was used to fix the embryo in place within the Petri dish and another one to scrap the yolk away and cut the tailbud. The thin layer of ectoderm was pulled off by needles to minimize the ectodermal contribution for the experiments and the AFM measurements conducted on the PSM surface. The developmental stage at dissection ranged from 5- to 8-somite stage.

For mechanical dissociation, three tails were collected in a microcentrifuge tube with 10 µL of L15 medium and mechanically dissociated by pipetting using a P20 pipette for 5 minutes. Dissociated cells were plated on PDMS microposts or glass-bottom dishes pre-coated with 0.2% F-127 Pluronic or 2% Matrigel for subsequent imaging. The mechanical dissociation process produced a mixture of single cells and cell aggregates, both of which were used for the analysis in this study.

For chemical dissociation, Accutase was diluted 1:1 with sterile PBS (50% Accutase) before use; as a guideline, ∼500 µL of 50% Accutase was used for four tailbuds. Tailbuds were incubated in 50% Accutase for up to 10 min. Following incubation, tissue suspensions were transferred carefully through sequential medium washes (total 1,000 µL medium for each four tailbuds across three wash droplets) to remove residual Accutase while minimizing premature dissociation during transfers. Tailbuds were transferred into a 2 mL low-retention tube (approximately 10 µL per tailbud; e.g., 40 µL for four tailbuds) and pipetted quickly for 20 times to separate into single cells thoroughly using wide-bore tips to reduce shear. Cells were plated into the prepared Grace chamber wells by adding 40 µL of the cell suspension per well. Mineral oil was overlaid on the ports to prevent medium evaporation during imaging, taking care to avoid overflow. Cells were allowed to settle for 30 min before imaging.

### Confocal time-lapse microscopy

Images were acquired using an inverted Olympus FV1200 confocal microscope equipped with a 20x objective (Olympus UCPlanFL 20x / 0.70 NA), PMT detectors, and a Z-direction compensation autofocus function. All chemical dissociation experiments were imaged on an upright Olympus FV3000 confocal microscope using either a 10× dry objective (UPLFLN10X2; NA 0.3; WD 10 mm) or a 20× water-immersion objective (XLUMPLFLN20XW; NA 1.0; WD 2 mm; field number 22 mm; no coverslip). Her1-Venus/Her7-Venus was excited using a 515 nm laser with 10% power (5% power on FV3000) and a scan speed of 12 μs/pixel (pixel dwell time) (10 μs/pixel on FV3000) and detected with a high-sensitivity GaAsP detector. Transmitted light images were captured using a transmitted light photomultiplier detector. The image size was 512 x 512 pixels, resulting in a resolution of 1.242 pixels/µm (0.805-5.632 pixels/µm for FV3000). Both transmitted light and YFP channels were imaged at 5-minute intervals for a minimum duration of 20 hours. The sample dish was maintained at 28°C using the Tokai Hit Stage Top Incubation System on FV1200, at ambient room temperature (approximately 20°C), without the Incubation System on FV3000. Multiposition scanning was configured to capture up to 14 positions per experiment.

### Cell selection and exclusion criteria

All isolated cells included in the analysis were viable, showed no signs of lysis or photobleaching during imaging, and survived for the full 10-hour tracking window. Dead cells indicated by no morphology dynamics or loss of motility for the last 10 frames within 10 hours, or the cells that could not to be tracked continuously (overlapping, moving out of frames, or touching other cells at the first frame) were excluded in the statistics. Oscillating cells were defined as those exhibiting ≥1 peaks in 10 hours in the mechanical dissociation data, and ≥1 peak in 20 hours in the chemical dissociation data. The different criterion for chemically dissociated cells was used because these experiments were performed at a lower temperature, resulting in slower oscillatory dynamics. In both cases, detected peaks were required to satisfy the specified prominence and minimum-separation values. To be noted, based on the time-lapse recordings, most PSM cells plated as single cells maintained a stable morphology without division, especially on softer substrates (0.6 and 2.9 kPa). On stiffer substrates (e.g., 1.2 MPa and Matrigel-coated glass), a higher proportion of cells exhibited migration and morphological elongation, but not necessarily increased division.

### Image analysis

Isolated cell Her1-Venus expression was tracked using Manual tracking with TrackMate^38^ in Fiji^39^. The tracked circle diameter was set to 10-15 µm to ensure coverage of the entire cell area across all frames. Peak detection and period statistics were obtained using a custom Matlab script with the *findpeaks* function, which smoothens the time series and identifies peaks based on local maxima, minimum period distance, and minimum prominence. The period was defined as the peak-to-peak time interval. For the criteria for period collection for single cells and aggregates generated by mechanical dissociation at 28 °C, the maximum period is defined at 145 min, approximately 2 times the mean on the control condition (Pluronic-coated glass). All tracks were isolated from the first frame, and the tracked cells could contact others but still be detectable within the cutoff. Oscillating cell percentage statistics were calculated by oscillating cells per all tracked cells using a 10-hour cutoff. Pie plots were used to show the percentage of all cell types from the first frame and tracked cells at 10 hours for each dataset (Figure S5).

The calculations of oscillating cell percentage were done across the counts from imaging positions. Positions with a sample size smaller than 5 were excluded from the final percentage estimates due to low statistics. Cell aggregates were defined as having at least 4 cells at the first frame and surviving for 10 hours. Oscillating cell aggregates were defined as having at least one oscillating cell, with peaks detected using the custom Matlab script within the 10-hour window. If cells split from the aggregates, all separated parts containing more than 4 cells were tracked. Cell aggregates were manually tracked using the Fiji/ImageJ plugin, Mastodon, with the tracked circle diameter set to approximately 4 µm larger than the object to minimize background noise impact on average intensity calculation. The algorithm smooths the time series and identifies peaks based on local maxima, minimum period distance, and minimum prominence.

### Automated image processing and analysis pipeline

Time-lapse microscopy datasets from chemical dissociation experiments were processed in Fiji/ImageJ using a standardized, macro-assisted pipeline to generate tracking-ready hyperstacks and TrackMate outputs for downstream quantification. First, raw microscopy files were split into brightfield (BF) and HER1-Venus channels using a custom split-channel macro (channelsplit_oif_to_bf_her1_.ijm). Separate input/output folders were assigned for the raw data, BF images, and HER1 images, while preserving the original raw-data filenames and appending channel-specific suffixes.

For object detection, BF images were segmented using the LabKit plugin. A classifier was trained on a representative dataset and then applied to the full dataset via the LabKit Batch Processor. Segmentation masks were saved into a dedicated “segmentation” output folder, and separate classifiers were trained for each surface type to account for condition-specific image features. The LabKit segmentation masks were then thresholded using a dedicated thresholding macro (thresholding_.ijm) to generate thresholded masks in a separate output folder.

Next, segmented and thresholded images were combined with the corresponding HER1-intensity images using a merge macro (merge_thresh_her1_.ijm) to generate final merged hyperstacks for tracking. The merge step used segmentation masks as input and exported merged hyperstacks to a designated output folder; merged file names additionally included the experiment date for traceability. Tracking was performed in TrackMate using representative merged hyperstacks for parameter training. A thresholding filter was applied in TrackMate, and separate TrackMate training files (.xml) were generated for single-cell versus aggregate datasets, including area-based filters to isolate single cells from aggregates; each training file was saved for reuse. For batch processing, all merged hyperstacks were processed using the corresponding single-cell or aggregate training .xml, and results were exported into condition-appropriate folders. The pipeline saved the TrackMate training file (.xml), AVI files for tracking verification, and CSV files for subsequent quantitative analysis. Consistent naming conventions and condition-specific folder organization (including surface and data type) were used throughout to maintain clear mapping between raw data, processed outputs, and analysis groups.

Downstream analysis was performed in MATLAB using a custom script Fin_single_cell_analysis.m, with separate file paths and output folders specified for single-cell and aggregate datasets. The script plotted frame/time versus HER1 intensity for each TrackMate track ID across CSV files, checked and sorted frame sequences for representative IDs, and identified oscillatory tracks by detecting peaks and troughs to calculate oscillation periods. The analysis then generated summary outputs including period-versus-time plots for oscillating cells, cycle-number plots for each condition, and oscillating-cell percentage comparisons across conditions.

### Fabrication of PDMS micropost arrays

Photolithography and deep reactive ion-etching (DRIE) techniques were used for the fabrication of the Si micropost mold. The PDMS micropost array was generated by replica molding^24,40^. PDMS prepolymer with a 10:1 base-to-curing agent ratio was poured into the Si micropost mold and cured at 110°C for 30 min. The negative PDMS template containing an array of holes was formed after peeling off from the Si micropost. Then the template was oxidized with oxygen plasma and passivated with trichloro (1H, 1H, 2H, 2H-perfluorooctyl) silane vapor overnight.

PDMS prepolymer with a 10:1 base-to-curing agent ratio was poured over the negative PDMS template, then covered by the cover glass (Fisher Scientific 12542B), and cured at 110°C overnight. The final PDMS micropost array was peeled from the negative PDMS template and subjected to sonication in 100% ethanol for 30 seconds, followed by dry-release with liquid CO_2_ using a critical point dryer (Samdri®-PVT-3D, Tousimis, Rockville, MD) to recover collapse of PDMS microposts during peeling process.

The array surface rigidities selected for PSM cell culture included 1.2 MPa (post diameter: 0.8 µm, post-to-post diameter: 1.6 µm, post height: 0.42 µm), 6 kPa (post diameter: 0.8 µm, post-to-post diameter: 1.6 µm, post height: 3.46 µm), 2.9 kPa (post diameter: 0.8 µm, post-to-post diameter: 1.6 µm, post height: 4.49 µm), and 0.6 kPa (post diameter: 0.8 µm, post-to-post diameter: 1.6 µm, post height: 7.57 µm). To attach cells to micropost tops, we functionalized Matrigel on the tops by contact printing. Firstly, PDMS stamps with a 30:1 base-to-curing agent ratio were generated and immersed in a solution containing Matrigel (2%; Corning) for 1 hour. Matrigel-coated PDMS stamps were then placed in contact with the PDMS micropost array pre-treated with UV-ozone (UV-ozone cleaner, Jelight, Irvine, CA) to transfer adhesive Matrigel from stamps to the tops of PDMS microposts. To avoid undesired cell adhesion to the side surfaces of microposts, PDMS micropost arrays were submerged sequentially in 100% ethanol (10 seconds), DI water (three times washing), and 0.2% w/v Pluronics® F-127 solution (Sigma-Aldrich; 30 minutes). Matrigel-coated PDMS micropost arrays could be stored in phosphate-buffered saline (PBS; Invitrogen) solution for up to a week before cell culture.

### Quantification of cell contractility of PSM cells

To quantify the traction forces exerted by isolated cells, we employed PDMS micropost arrays. The PDMS microposts beneath the isolated cells were stained with Fibrinogen, Alexa Fluor™ 647 Conjugate (Invitrogen™) and imaged using an inverted Olympus FV1200 confocal microscope equipped with an Olympus UPlanSApo 40x 1.25 Sil objective. Time-lapse images were analyzed using a custom-developed MATLAB script^24,40^. The script fitted the deviation of each post’s centroid from its original position, determined by the free and undeflected posts. The horizontal traction force was then calculated by multiplying the post centroid deviation by the nominal spring constant K, which was generated through finite element model (FEM) simulations^24,40,41^.

### Oscillator model

We modeled the her1 genetic oscillator based on a time-delayed negative feedback model adapted from Negrete et al.^28^, which is described by the delay differential equation (DDE):

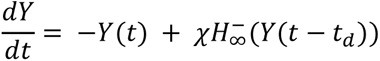

Here *Y* is the Her1 protein concentration, 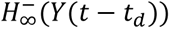 is the delayed negative feedback with an explicit time delay *t*_*d*_, and *χ* is the Her1 production rate. 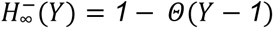, where *Θ* is the Heaviside step function.

Time is measured in units of the Her1 protein half-life *T*_0_ = 5 min. The feedback delay was set to *t*_*d*_ = 7 (dimensionless), corresponding to a physical delay time of *T*_*d*_ = *t*_*d*_ × *T*_0_ = 35 min, which accounts for the combined transcription and translation delays and reproduces the experimentally observed oscillation periods of 60-100 min in isolated PSM cells in Figure 2D, longer than the ∼30 min zebrafish embryonic segmentation clock period, consistent with slowed kinetics in dissociated cultured cells.

For a free-running oscillator without mechanical input, *χ* is constant. This system exhibits a sharp oscillation threshold at *χ* = 1: for *χ* < 1 the system decays to a fixed point and no oscillations occur; for *χ* > 1 sustained oscillations emerge.

It has been observed in similar contexts that mechanical forces, acting through the YAP pathway, can create a thresholding effect for the onset of oscillations, and that Notch can help rescue these oscillations^11^. With these prior results and our experimental findings in this study, and to incorporate the effects of substrate rigidity, we treat *χ* as a function of both Notch and Yap activity, such that *χ* = *f*(*Notch*)*g*(*YAP*(*S*)) where *f* is a monotonically increasing function of Notch (activator) and *g* is a monotonically decreasing function of YAP (inhibitor). Additionally, we modeled YAP following a Hill function relationship with substrate rigidity *S*:

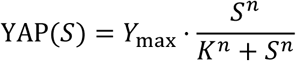

consistent with the experimental observation of switch-like YAP translocation in response to increasing rigidity^20^. The mean production rate then follows:

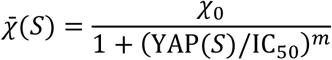

where *χ*_0_ is the production rate in the absence of YAP inhibition and IC_50_ is the YAP level at half-maximal inhibition (*m* = 2 fixed). Specifically, for the results in Figure 2, Notch input was held constant f(Notch)= 1.

To capture cell-to-cell variability, *χ* was sampled from a Gaussian distribution, with mean *χ*ˉ(*S*) and standard deviation *σ* = *σ*_frac_ ⋅ *χ̅*(*S*), where *σ*_frac_ is a fitted parameter. All six model parameters (*χ*_0_, *K*, *n*, IC_50_, *σ*_frac_, *Y*_max_) were fit by least-squares to the experimentally measured fraction of oscillating cells across substrates, with 95% confidence intervals estimated by bootstrap resampling (*n* = 1000). The fitted parameter values are reported in Supplementary Table S4. The fitted *σ*_frac_ indicates that cell-to-cell variability in *χ* is large, consistent with the broad distributions of oscillatory behavior observed experimentally. Individual simulated trajectories were integrated using MATLAB’s dde23 solver, and oscillations and periods were extracted from each trajectory (Figure S6A). Fitting this model to the experimentally measured fraction of oscillating cells across substrates yielded a rigidity threshold of 3.77 kPa (95% CI: 2.58–5.37 kPa) (Figure 2F), consistent with the experimentally observed transition range of 2.9-6 kPa (Figure 2B) and the reported switch-like YAP nuclear translocation threshold of ∼5 kPa in cultured mouse embryonic fibroblasts^20^ (Figure S6B-C).

This model naturally accounts for the stiffness-dependent decline in the fraction of oscillating cells: as rigidity increases, YAP activity rises, suppressing *χ*ˉ and driving more cells below the oscillation threshold *χ* = 1.

The model also provides a mechanistic basis for the enhanced oscillatory fraction in cell aggregates: Notch signaling between cells increases *f*(Notch), which increases *χ* above threshold in a larger fraction of cells, consistent with our experimental results from the cell-aggregated data.

### Mean squared displacement

The mean squared displacement (MSD) gives a measure for the type of motion displayed by particles in a given time interval^42^. For each stage position, a background fixed point was tracked to account for slide movements from the stage. These displacements were subtracted from the cell tracking data in that position. Windowed MSD plots were generated for oscillating and non-oscillating cells on varying rigidity. The equation *MSD* (*t*) = < *r*^2^ > = < (|*r*(*t* + *to*) − *r*(*to*)|^2^ > and time from 0 to 600 min was used to produce plots. A windowed MSD calculation was also generated for each individual cell to generate span plots using the equation *MSD* (*Tau*) = < (|*r*(*t* + *Tau*) − *r*(*t*)|^2^ >. We selected all time frames and Tau values to generate smooth MSD estimates for each cell. The windowed MSD was verified with two separate algorithms and MSD at to = 0 was compared to the windowed MSD. Span plot areas were colored by 10% quantiles in MSD data.

Diffusion coefficients (D) were calculated using the equation MSD=2pDt where p=2 is the number of dimensions. We performed a linear least squares fit centered at the origin for each individual cell displacement track to evaluate diffusion in each condition. Span plot areas were colored by 10% quantiles in MSD data.

### Circularity

Isolated cell circularity was collected manually by tracing the boundaries of cells in Fiji (Image J) software. Circularity was collected for 120 frames over 10 hours for the selected cells. The degree of circularity was calculated using the *equation 4π*(*Area*/*Perimeter*^2^).

Time to shape transition was calculated as follows: a time-lapse was divided into 40-minute windows, a given frame for non-oscillating cells was considered to have undergone a significant shape transition if it’s 5-frame moving average circularity was 2 standard deviations less than the moving average of the oscillating cells’ circularity; If an entire 40-minute window consisted of frames designated as having undergone a shape transition, the first frame was marked as the time to shape transition for the condition.

### Verteporfin treatment for YAP inhibition

To inhibit YAP activity, a Verteporfin treatment solution was prepared in E3 medium at 200 μM with 1% Dimethyl sulfoxide (DMSO) from a 20 mM Verteporfin stock dissolved in 100% DMSO. A vehicle control solution containing 1% DMSO in E3 was prepared in parallel. Embryos were transferred into either 200 μM Verteporfin in 1% DMSO, 1% DMSO alone, or untreated E3 at the 3-somite stage and maintained in the corresponding condition until screening. Before tailbud dissection, embryos were rinsed three times with E3. Tailbuds were then dissected and chemically dissociated as described above in Fish husbandry and dissociation for zebrafish tailbud cells. Dissociated cells were plated into Grace Bio-Labs CoverWell perfusion chambers coated with either 0.2% F-127 Pluronic or 2% Matrigel, overlaid with mineral oil to prevent evaporation, and allowed to settle for 30 minutes before imaging. Oscillation percentage, oscillation cycle number, and clock period were quantified for both single cells and cell aggregates under untreated, DMSO-only, and Verteporfin-treated conditions.

### Immunofluorescence staining

Zebrafish tailbud cells were chemically dissociated from the same posterior regions used for oscillation analysis and plated on Nunc™ Glass Bottom Dishes pre-coated with F-127 Pluronic or 2% Matrigel. Cells were allowed to reaggregate and were cultured at 28°C for 5 −7 hours, which was the minimum time required for most cells to adhere to the substrate. Cells were then fixed with 4% paraformaldehyde (PFA) in Milli-Q water for 15 minutes at 20℃, rinsed with PBS, permeabilized with 0.1% Triton™ X-100 for 10 minutes at room temperature, and blocked with 0.1% Bovine Serum Albumin (BSA) in PBS for 30 minutes at room temperature.

To distinguish cell types and fates, samples were incubated with anti-SOX2 antibody diluted 1:200 in 0.1% BSA for 2 hours at room temperature, followed by two PBS washes. From this step onward, samples were protected from light. Samples were then incubated with Alexa Fluor 488-conjugated secondary antibody diluted 1:400 in PBS for 2 hours at room temperature, followed by two PBS washes. Samples were then blocked again with 0.1% BSA in PBS for 30 minutes at room temperature and incubated with anti-Brachyury antibody diluted 1:200 in 0.1% BSA for 2 hours at room temperature, followed by two PBS washes. Samples were then incubated with Alexa Fluor Plus 405-conjugated secondary antibody diluted 1:400 in PBS for 2 hours at room temperature, followed by two PBS washes. After an additional blocking step, samples were incubated with anti-VegT antibody, which was received pre-diluted and used directly without further dilution, for 2 hours at room temperature, followed by two PBS washes. Samples were then labeled with the Alexa Fluor 594 Zenon™ Mouse IgG2a Labeling Kit at room temperature according to the manufacturer’s instructions.

To quantify YAP, samples were incubated with anti-YAP1 polyclonal antibody diluted 1:250 in 0.1% BSA for 2 hours at room temperature, followed by two PBS washes. From this step onward, samples were protected from light. Samples were then incubated with Alexa Fluor Plus 405 - conjugated secondary antibody diluted 1:500 in PBS overnight at 4℃, followed by two PBS washes.

Images were acquired using an Olympus FV3000 confocal microscope equipped with a 20x objective at 512 × 512 pixel resolution with 1×, 2×, or 3× digital zoom.

## Data Availability

Source Data are provided with this paper. Raw microscopy imaging datasets are available from the corresponding author upon reasonable request due to file size and data-management constraints.

## Code Availability

Custom MATLAB analysis scripts used in this study are available at the Yang Lab GitHub repository: https://github.com/YangLab-um/her1-rigidity

